# *Ceudovitox:* a novel pseudotyped-virus screening platform to identify cell entry factors for high consequence infections

**DOI:** 10.1101/2025.07.16.665096

**Authors:** Tom Stiff, Elle A Campbell, Salih Bayraktar, Christine Reitmaye, Edward Wright, Leandro Castellano

## Abstract

Development of scalable and highly adaptable platforms to characterise high consequence infections are essential for limiting the impact of future viruses with pandemic potential. As part of this, an understanding of the virus-host interactions is vital, with the cell entry mechanism being crucial for the development of therapeutics and vaccines. Current approaches in this field depend on assays with authentic (live) viruses that require high containment facilities. However, these are hindered by high costs, need for highly trained staff, slow processing times and limited scalability. Using pseudotyped viruses (PV) expressing the chikungunya (CHIKV) envelope protein as a proof-of-principle, we developed a novel screening platform, *Ceudovitox*, to identify cellular factors involved in viral entry. PVs were engineered to express the herpes simplex virus-1 thymidine kinase, which following addition of ganciclovir, can selectively kill infected cells. Then a heterogenous pool of knockout cells were produced using the CRISPR-Cas9 library. Infection of these cells with the “killer” PV system permitted positive selection of cells refractory to viral infection and, through next generation sequencing, identification of a number of factors involved in CHIKV entry. Matrix metalloproteinases were identified as novel entry factors and demonstrated that MMP-targeting drugs efficiently inhibit PV and authentic CHIKV infections. These results suggest this platform holds great promise as a pandemic-preparedness tool that increases the capacity and speed of screening for cell entry factors. It is also a safe platform able to dissect the gene network involved in virus entry. *Ceudovitox* will help identify new therapeutic targets, thereby aiding the development of treatments against future outbreaks or pandemic pathogens and making a significant contribution to the “100 days” mission.

**Author Summary:** The increased threat of the emergence of new viral pathogens with pandemic potential highlights the importance of developing scalable high-throughput screening platforms to quickly characterise new emergent viral pathogens. Improving understanding of virus-host interactions, including how viruses enter cells, can help fast development of new targeted therapeutics and reduce pandemic burdens. Current approaches rely on using ‘live’ viruses, which are highly infectious and must be manipulated in high containment facilities, which are slow and cumbersome. Our new screening platform, *Ceudovitox*, is a safer, faster and more scalable alternative. Through using pseudotyped viruses, which can only undergo one replication cycle, we are able to distinguish virus entry factors of high containment viruses within highly abundant low containment facilities, allowing us to quickly characterise potential entry factors of high containment viruses. Within this study, *Ceudovitox* was shown to recognise both known and novel entry factors of Chikungunya virus; a virus which causes chronic arthralgia and arthritis. Matrix metalloproteinases were amongst novel entry factors identified and were seen to reduce virus infection in both pseudotyped and authentic virus assays, demonstrating how *Ceudovitox* can correctly discover virus entry factors of ‘live’ high containment viruses. *Ceudovitox’s* utility as a screening platform for high containment viruses and new emerging pathogens, has the potential increase understanding and speed up the development of new therapeutics during pandemics, helping to reach the ‘100 Day Mission’ of pandemic preparedness.

## Introduction

The current human population has created the perfect environment to favour emergence and further transmission of infectious viruses, resulting in a number of viral pandemics throughout the 20^th^ and 21^st^ centuries [1]. Pandemics have also been predicted to become 38% more frequent during peoples’ lifetimes [2]. Therefore, the importance of the development of fast, flexible and scalable workflows, which can support identification of therapeutic targets of numerous emerging pathogens, is an international priority. Innovation of adaptable platforms, that can be used in highly abundant containment level 2 laboratories, are therefore fundamental to meeting these international initiatives for pandemic preparedness. This includes the “100 days” mission to develop and deploy new therapeutics and vaccines for new emerging pathogens within 100 days of the first reported case [3].

A number of pathogens have been highlighted by the World Health Organization (WHO) as priority pathogens for epidemic and pandemic research preparedness, as they have no licensed vaccines or antivirals. For example, chikungunya virus (CHIKV), which is known to cause chronic arthritis in infected individuals for as long as 3 years after initial infection in recent outbreaks, yet has no licensed antivirals and patients are currently treated by unspecific supportive care and anti-inflammatory drugs [4, 5]. Many of these viruses are highly infectious and therefore, research involving the authentic pathogens can only occur in high containment facilities (*e*.*g*.: containment Level 3 and 4 laboratories). Work within these facilities is cumbersome and slow and can only be undertaken by highly trained laboratory workers. This hampers efforts to improve our understanding of these viruses, including their interaction with infected cells and their hosts, which underpins the identification of new therapeutic targets and vaccine design.

CHIKV is a hazard group 3 pathogen, classified within the genus *Alphaviridae* and the family *Togaviridae*. CHIKV is personified by chronic joint and muscle pain in infected individuals. Symptoms can sometimes last for months to years, reducing patients’ quality of life and their ability to work, leading to subsequent economic implications. CHIKV outbreaks used to be primarily restricted to Asian and African countries, but now the virus is reported to be prevalent in over 60 countries worldwide, causing this virus to be international concern [4]. However, CHIKV currently has no licensed antivirals, and its lifecycle is far from fully understood. For example, Mxra8 has recently been identified as a cellular receptor for CHIKV [6, 7]. However, CHIKV is capable of infecting cells lacking Mxra8, suggesting Mxra8 is a minor factor and more importantly host viral entry factors are yet to be identified, which could act as future therapeutic targets [6]. Therefore, highlighting the need for undertaking functional screening approaches able to identify the gene network responsible for CHIKV entry. Ideally this research needs to be expanded to lower containment facilities to increase throughput, using scientific tools such as replication-deficient models of viral infections, including viral-like particles and pseudotyped viruses (PVs) [8, 9].

PVs’ chimeric nature restricts them to only one infection cycle, making PVs a safer alternative to handling authentic, replication-competent viruses. PV can be used for various studies, such as the detection of neutralising antibodies from serological samples and the study of viral entry. A meta-analysis found a high level of correlation between neutralisation studies completed with live viruses and PVs [10]. Likewise, the development of *Arenavirus* pseudotyped VSV systems has allowed viral entry pathways to be deciphered for these neglected pathogens, showing that PV are an appropriate surrogate for studying the early events in the virus lifecycle [11]. Many PVs also encode luminescent reporter genes, allowing easy and fast identification of PV infectious titres. Therefore, PVs are an effective tool for developing quick and extensive screening protocols for therapeutic and vaccine target identification within low containment facilities.

In this study, our novel screening system, *Ceudovitox*, utilises a CHIKV lentiviral PV encoding the HSV-1 thymidine kinase (HSV1-TK), which is cytotoxic in the presence of ganciclovir (GCV) and when combined with our PV can kill infected cells with a >90% efficiency. Utilisation of this PV system in the context of a heterogenous population of cells, all deficient in one singular gene, allowing for the positive selection and retention of cells refractory to viral infection. This phenotype is acquired by using Cas-9 to specifically knock-out genes important for host cell entry, which can ultimately be identified by deep sequencing of the refractory cells.

*Ceudovitox* overcomes the hurdles encountered by existing systems used to study viral entry. In addition to the superior scalability, applicability and adaptability of our system, the successful engineering of our PV to kill infected cells means we can study viruses in highly physiologically relevant cells that might not be possible with authentic viruses due to the lack of cytopathic effect. Additionally, utilisation of PVs, which are replication defective, means our assay can specifically focus on virus binding and entry factors, unlike authentic virus assays. Finally, *Ceudovitox* can be applied to viruses where no isolate has been obtained or critical reagents required for authentic virus assays, such as antibodies for staining, are not available. These points highlight the considerable benefits of this approach as a fast, scalable screening platform for identifying therapeutic targets for high consequence viruses.

## Results

### Selective cell killing using TK CHIKV-PV and ganciclovir

First, to mirror CHIKV entry dynamics a replication-incompetent pseudotyped virus (PV) was designed to bear the CHIKV envelope proteins (E3-E2-E1), which are solely responsible for CHIKV attachment and subsequent entry into infected cells [12]. Along with direct targeting cells that would be subjective to CHIKV infection, the PVs also expressed the herpes simplex virus-1 thymidine kinase (HSV1-TK) gene (Figure 1A). HSV1-TK can selectively kill transduced cells, due to the production of cytotoxic GCV-triphosphate by HSV-TK after subsequent ganciclovir treatment [13]. Supporting this, increased concentrations of transduced HSV1-TK caused a direct increase in cell death in HEK293T/17 and Huh7 cells (Figure 1B). Therefore, the engineered PV could act as a strong selective pressure to easily distinguish between cells resistant or susceptible to CHIKV infection within a heterogenous cell population generated from CRISPR knockout systems.

**Figure 1:**
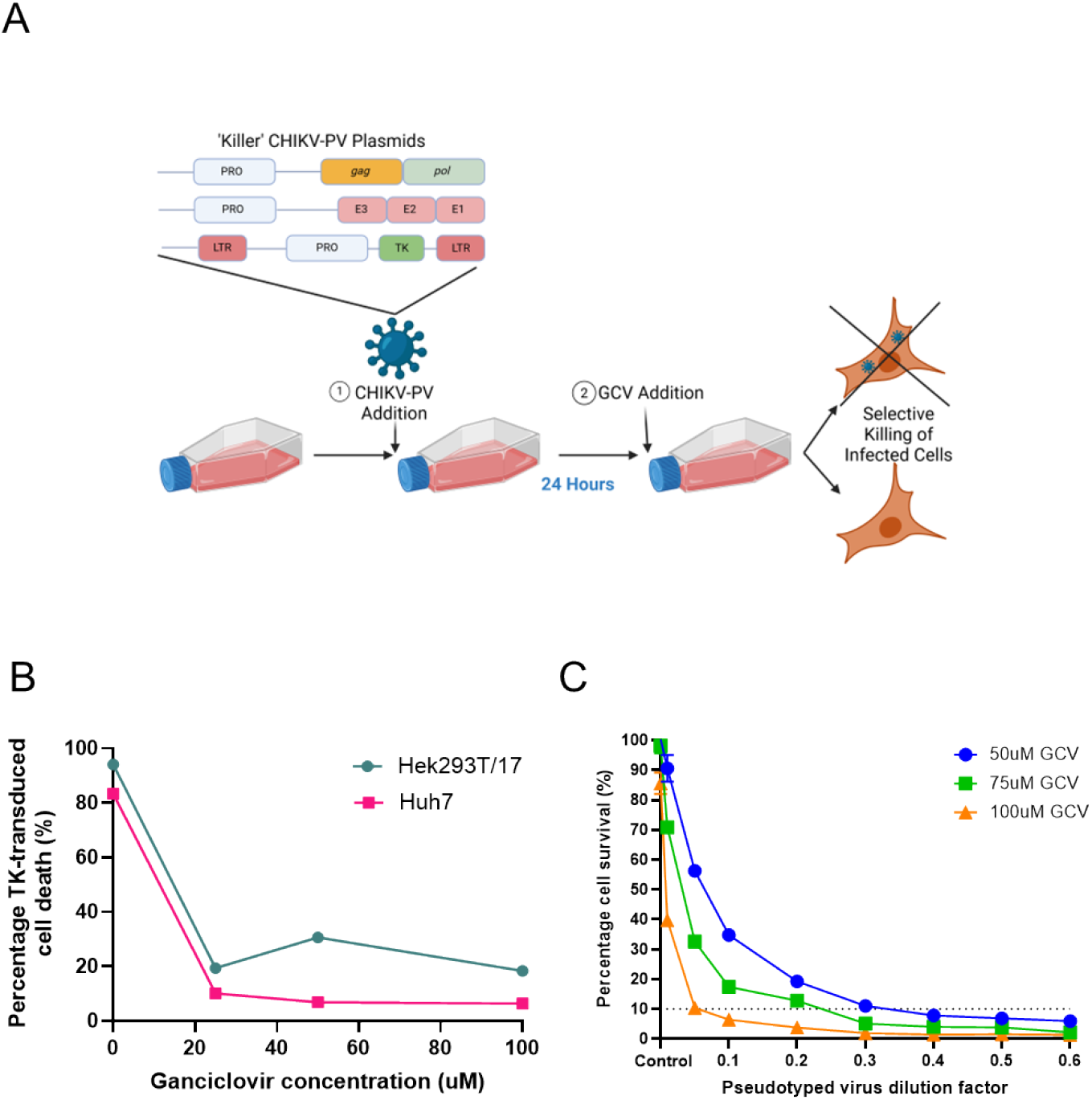
Infection of cells with a PV encoding the HSV-1 TK gene, followed by subsequent treatment with ganciclovir, allows selective killing of infected cells. (A) PVs encoding the HSV-1 TK and Chikungunya virus envelope protein were generated using a three-plasmid system. A549 cells were infected with the ‘killer’ PV and 24 hours later treated with GCV, causing selective killing of virus infected cells. Designed with Biorender. Created in BioRender. Castellano, L. (2025) https://BioRender.com/e25c8u2 (B) Percentage cell viability in HSV-1 thymine kinase transduced cells compared to non-transduced control in HEK293T/17 (blue line) and Huh7 (pink line) cells. (C) Virus titres and GCV concentrations were optimised prior to GeCKO screening experiments to allow high percentage of infected/killed cells. Different GCV concentrations shown - 50 uM (blue line), 75 uM (green line) and 100 uM (orange line).

Therefore, this system was optimised to determine the titre of virus required to kill 90% of cells after GCV addition to function as a strong selective pressure in GeCKO CRISPR knockout screen, with a goal of finding novel factors involved in CHIKV entry. TK CHIKV-PV was titrated onto the target cells in doubling dilutions and GCV added to the cells 24 hours later. A concentration of 75 nM GCV effectively kill over 90% of cells after 48 hours treatment (Figure 1C). Therefore, this concentration of GCV was utilised within the subsequent GeCKO screen.

### Infection of a pool of differential single-gene knockout cells allows identification of novel CHIKV entry factors

The ‘killer’ PV system was challenged against a population of cells previously treated with the GeCKO lentivirus library [14]. This library contains over 123,411 sgRNAs sequences allowing targeted knockout of over 19,050 genes [15]. This library can therefore be used to produce a heterogenous pool of cells, with each cell missing the expression of a different singular gene. If one of these genes were involved in virus entry and was subsequently knocked out of the genome, the virus would no longer be able to infect the cell. Therefore, transforming a once permissive cell to have a restrictive phenotype. Therefore, infecting these knockout cells with the ‘Killer’ TK CHIKV-PV allowed positive selection of cells resistant to CHIKV entry. While susceptible cells were killed through GCV treatment, surviving resistant cells multiplied and become dominant within the population. This enrichment was then quantified by analysing next generation sequencing data of the sgRNAs incorporated in the genomes and their differential identification in both infected and control GeCKO population cells (Figure 2A.) Mapping of these enriched sgRNA with their target’s genes allowed identification of candidate genes with an involvement in CHIKV entry dynamics [16].

**Figure 2:**
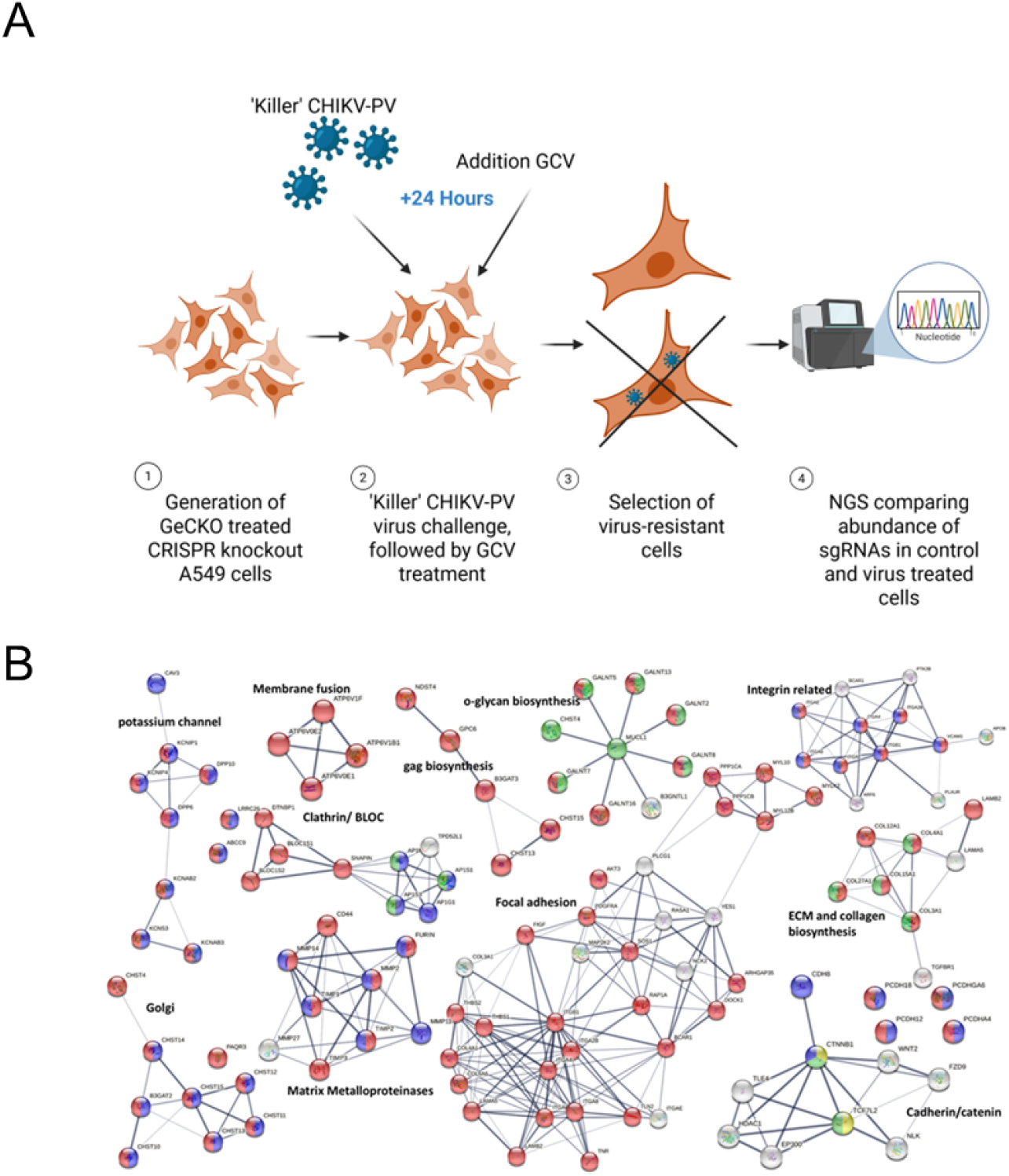
Infection of GeCKO knockout cells with ‘killer’ PV expressing the CHIKV envelope proteins allows identification of genes related to CHIKV entry. (A) A549 cells were treated with the GeCKO library encoding 19,050 sgRNAs targeting over 19,050 genome coded genes, producing a heterogenous A549 CRISPR knockout cell population. Cells were then infected with ‘killer’ TK CHIKV envelope encoding PVs and 24 hours treated with ganciclovir. This allowed selection and retention of infection-resistant cells, who no longer contained CHIKV entry factors. Cells were then allowed to grow for 14 days, allowing enrichment of infection-resistant cells and their sgRNAs targeting virus entry factors. Cells were then harvested and subjected to next generation sequencing to detect differences in sgRNA abundance in control and infected cells. Created in BioRender. Castellano, L. (2025) https://BioRender.com/ossvr9q (B) MAGeCK analysis identified a number of genes whose sgRNAs were enriched within the infection-resistant cells. A cutoff of a positively p-value produced by the MAGeCK RRA of <0.05 and an FDR <0.25 was used to find significant hits. STRING analysis was then used to identifying common pathways between positive selection hits. Both known and novel pathways were identified to be involved in CHIKV entry, including MMPs.

Genes essential for CHIKV virus entry were then identified by looking at genes whose sgRNA were significantly positively selected for during the screen, using significant cutoffs of a p-value of <0.05 and an FDR of <0.25 calculated by the MaGECK robust rank aggregation algorithm (RRA) [16]. Enriched genes were found to be involved in several different cellular pathways, according to a STRING analysis [17]. Interestingly, matrix metalloproteinases (MMPs), which have both previously not been recognised as CHIKV entry factors, were detected as important cellular pathway involved in CHIKV entry and were considered for further investigation (Figure 2B).

### Broad-spectrum MMP drugs inhibit CHIKV PV and authentic CHIKV infection

As the screen had identified MMPs as potential novel inhibitors of CHIKV infection, a number of broad-spectrum MMP drugs were tested for their efficiency of preventing CHIKV-Envelope PV virus (Figure 3A). Unlike the ‘killer’ PV virus used in our initial screen, in place of the thymine kinase gene PV used in this assay encoded the luciferase gene, whose luminescent signal could be used as a direct quantification of viral infection. Broad spectrum MMP inhibitors, marimastat and GM6001, and previously published CHIKV entry inhibitors (arbidol) consistently inhibited CHIKV PV entry [18-20]. All MMP were non-cytotoxic at concentrations of <50 nM, with percentage healthy vs vehicle above 100% (S1 Appendix).

**Figure 3:**
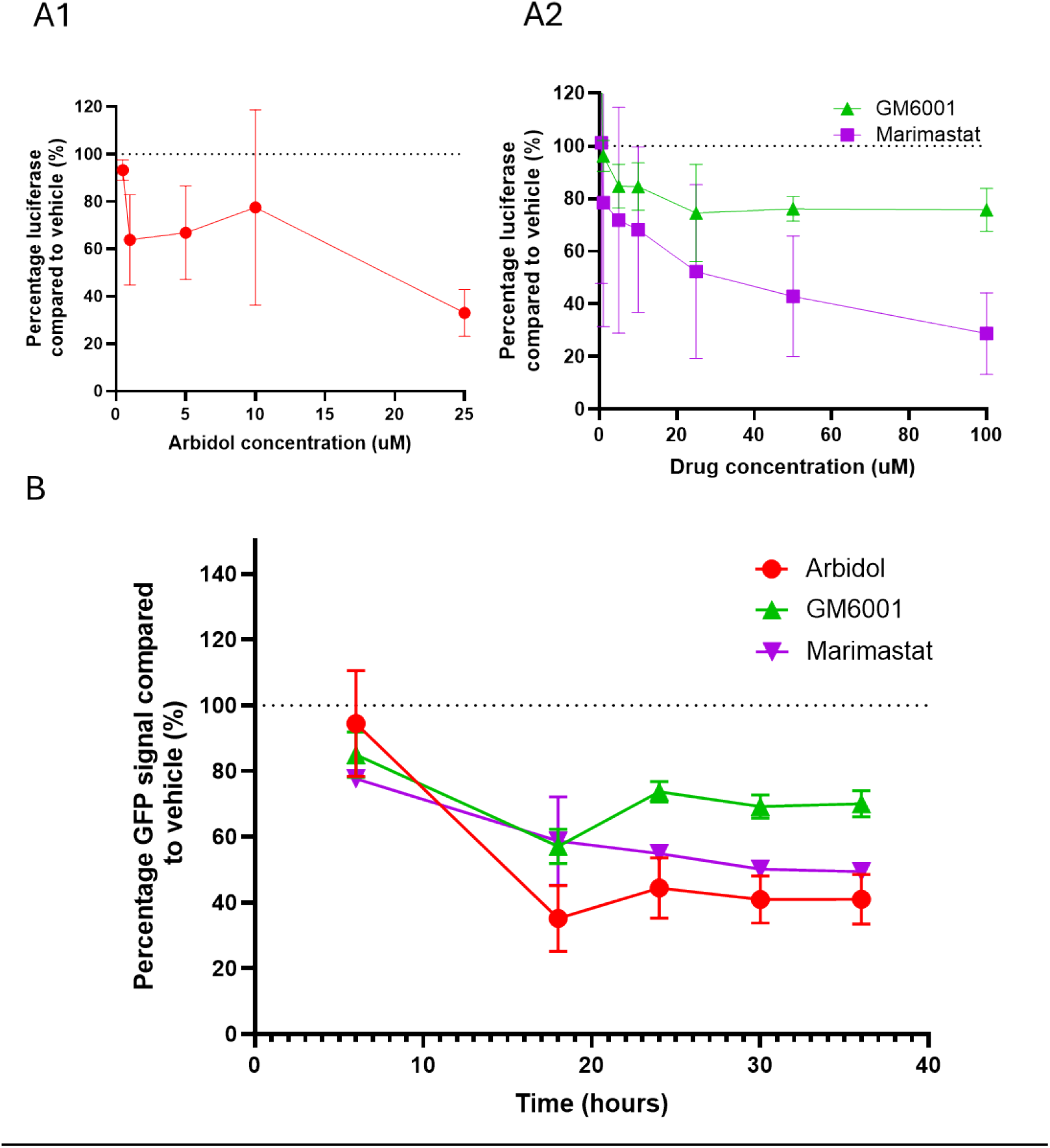
Inhibition of matrix metalloproteinases (MMP) inhibits Chikungunya infection. (A1) Known Inhibitors, arbidol (red line)), of Chikungunya entry and (A2) broad-acting MMP inhibitors, Marimastat (purple line) and GM6001(green line), were found to inhibited CHIKV-PV entry. (B) Treatment with Matrix Metalloproteinases (MMP) inhibiting drugs also reduces infection rates of live Chikungunya-GFP virus. In Figure 4B the following drugs are shown as Arbidol 10 uM (red line), GM6001 10uM (green line) and marimastat 50 uM (purple line).

To validate the eligibility of targeting MMPs to reduce CHIKV viral entry, marimastat and GM6001 were tested in authentic virus-based inhibition assays using a CHIKV virus expressing the GFP protein (Figure 4B). These assays showed similar dynamics to CHIKV PV assays, demonstrating CHIKV PVs applicability as a virus entry model and validating the credibility of protein target hits identified through the initial *Ceudovidox* PV screen. In live virus assays, a concentration of 50 uM of marimastat was found to reduce GFP signal by 50 % at 18 hours post infection, again highlighting these proteins applicability as therapeutic targets for CHIKV infection (Figure 3B).

**Figure 4:**
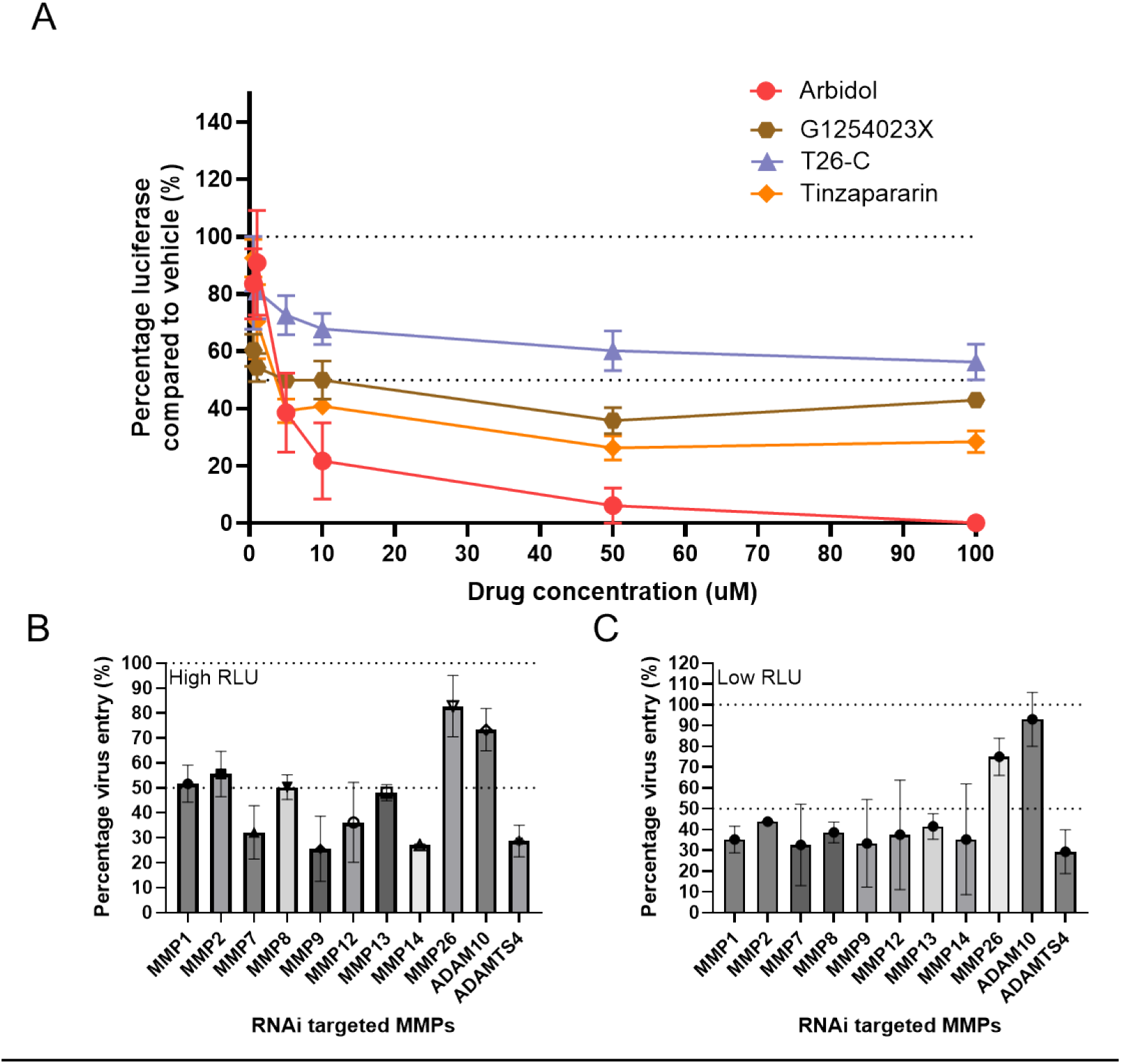
Inhibition of specific MMPs inhibits CHIKV-Envelope PV entry. (A) A panel of MMP inhibiting drugs targeting specific MMP related proteins were screened for their efficiency at preventing CHIKV-Envelope PV entry. These drugs targeted ADAM10 (G1254023X, brown line), MMP13 (T26-C, purple line) and ADAM54 (Tinzapararin, orange line). Arbidol (red line) was used as positive control. (B&C) RNA interference was performed on a range of MMP related proteins to selectively silence their expression and demonstrate its knock-on effect on cells susceptibility to CHIKV-Envelope PV entry at High (B) and Low (C) RLU.

### Inhibition of specific MMP and MMP-related proteins restrict CHIKV-Envelope PV entry

To further understand what MMP proteins were involved in CHIKV entry dynamics, a number of drugs targeting singular MMP or related proteins containing a disintegrin and also a metalloproteinase domain (ADAM) were screened for their efficiency at reducing CHIKV entry (Figure 4A). All tested drugs reduced viral entry by 50% or below, with no negative effects on cell viability (Figure 4A and S1C Appendix). Hence, these results show that inhibition of multiple MMP and MMP-related proteins independently can restrict CHIKV entry (Figure 4A). These results were later supported through performing specific gene expression silencing of multiple MMP related proteins through RNA interference (Figure 4B). Multiple MMP proteins were again shown to reduce viral entry by over 50%, suggesting an involvement of this viral family in CHIKV entry dynamics (Figure 4B).

## Discussion

*Ceudovitox* is a novel screening platform allowing identification of candidate cellular genes involved in virus entry. This screen employs the use of a replication-incompetent PV system, allowing cell entry of high consequence viral pathogens to be deciphered in widely available low containment facilities. As a proof of concept, we have demonstrated how producing PVs encoding the cytotoxic HSV-TK gene and bearing the CHIKV envelope protein can allow selective cell killing. Challenging these systems against a population of CRISPR knockout cells, all missing different cellular genes, allowed identification of known and novel CHIKV entry factors, which were later validated with live virus assays. Therefore, demonstrating how this system can be used to recognise new therapeutic targets and decipher viral entry dynamics of new emerging pathogens.

Our screening method utilised the transduction of A549 cells with guide RNAs derived from the CRISPR-Cas9 knockout (GeCKO) library [15]. This has been shown as a powerful tool to recognise key host factors involved in several virus life cycles [21-23]. Unlike RNA interference (RNAi) approaches, which target mRNA transcripts and knockdown subsequent protein expression, GeCKO guide RNAs directly target the genomic DNA. This allows production of indel mutations and complete knockout of target genomic elements across the genome, including introns, promoters and inter-genic regions [15]. Therefore, allowing a more comprehensive understanding of host entry factors on virus life cycle dynamics.

Furthermore, unlike live viruses, PVs are unable to replicate and egress from infected cells, allowing virus entry to be clearly and singularly analysed [24]. Expression of the HSV-TK gene allows selective killing of infected cells with over >90% efficiency (Figure 1B and 1C). Therefore, this system could also be used to identify host factors for non-cytopathic viruses, unable to cause cell death after infection [25]. Likewise, the envelope protein are the main determinants of receptor binding dynamics for many high consequence viral infections and these PV systems can be easily adapted to contain alternative virus envelope proteins of emerging infections [26].

A limitation of the screen includes the presence of false positives occurring through off-target effects generated by the chimeric nature of the PV virus or incomplete cell killing by GCV. However, further optimisation including multiple rounds of infection and challenging knockout cells with PVs comprising different cores bearing the CHIKV envelope protein would reduce these effects.

Despite this, our screen was seen to successfully identify known and novel CHIKV entry factors. Glycosaminoglycans (GAGs) biosynthesis proteins were found to be involved in CHIKV entry through a STRING analysis, derived from looking at the top screen hits (Figure 2B). GAGs have been previously described to be involved in entry process of a number of viral pathogens, including CHIKV. GAGs, predominantly heparan sulfate, directly interact with CHIKVs envelope 2 protein and act as an attachment factor for multiple CHIKV strains [27, 28]. Attachment factors allow longer retention of viral particles on targeted cells, increasing their chance of interacting with receptors capable of endocytosis [29]. Therefore, demonstrating that our screen can identify host factors involved in authentic virus entry.

However much more interestingly, the STRING analysis also identified novel factors involved in CHIKV entry factors (Figure 2B). MMPs are a group of enzymes known to have endopeptidase activity and have a heavy involvement in degradation of the extracellular matrix, which is known to be a mechanical barrier to infection [30, 31]. Collagen and ECM biosynthesis enzymes were also highlighted in the STRING analysis, again supporting MMPs possible role in CHIKV virus entry dynamics (Figure 3). We found that treatment of cells with known MMP inhibitory drugs including Marimastat and GM6001 inhibited entry of PV infection (Figure 3A). Likewise, the same effect was mirrored in live CHIKV virus assays, with both drugs seen to reduce infection of a GFP-encoding CHIKV virus (Figure 3B). Direct knockdown of multiple MMPs were seen to be reduce viral entry in PV assays. (Figure 4). MMPs are also seen to be implicated in CHIKV infection in mosquitoes, with higher MMP expression increasing CHIKV transmission efficiency through increased permeability of the extracellular matrix basal lamina lining of *Aedes aegypti* midguts [32]. Therefore, further work deciphering MMP involvement in CHIKV infection in both human and mosquito hosts will support both therapeutic and vector transmission control strategies.

Overall, this study demonstrated how use of PV viruses encoding both the envelope genes of a virus of interest and the HSV-TK gene can (1) allow selective killing of infected cells, (2) leading to recognition of viral entry genes, which can be (3) validated by live authentic virus assays. This screening platform therefore strengthens global preparations against epidemics and pandemics, addressing crucial gaps in the field and setting new standards for global health innovation.

## Methods

### Cell lines

A549 cells (ATCC Number: CCL-185) were grown in DMEM (Sigma) supplemented with 10% FCS, 100 U/ml penicillin, and 100 mg/ml streptomycin. HEK293T/17 cells (ATCC Number: CRL-11268) were grown in DMEM (Sigma) supplemented with 10% FBS and maintained at 37^°^C with 5% CO_2_. BHK-21 baby hamster kidney cells (CLL-10, ATCC), they were cultured in Glasgow’s minimal essential medium (Life Technologies), supplemented with 10% heat-inactivated FBS and 10% TPB and 100 U/mL penicillin, and 0.1 mg/mL streptomycin at 37 °C and 5% CO2.

### Plasmids and primers

For development of the GeCKO library, the lentiviral construct used for the GeCKO library was lentiCRISPRv2 (Addgene) with psPAX2 (Addgene) and pMD2.G (Addgene) as VSV-G envelope-expressing plasmids used with lentiviral vectors to produce lentiviruses.

In the case of producing the PV to enable selective pressure during the screening assays, pCAGGS CHIKV (S27) E was generated by cloning the E3-E2-E1 cassette (Accession: AF369024.2) into pCAGGS so its expression was driven by the chicken β-actin promotor. The TK gene was PCR amplified from pAL119-TK (Addgene, RRID:Addgene 21911) and cloned into pCSFLW in place of the firefly luciferase gene, to produce pCSTKW [33]. pAL119-TK was a gift from Maria Castro (Addgene plasmid # 21911 ; http://n2t.net/addgene:21911 ; RRID:Addgene_21911). Successful cloning was confirmed by Sanger sequencing.

### Authentic (live) CHIKV preparation

All live virus experiments with CHIKV were performed under Containment Level 3 (CL-3) conditions. CHIKV (isolate LR2006OPY1) harbouring a ZsGreen marker and a duplicated subgenomic promoter in the intergenic region was rescued from DNA plasmids containing a SP6 promoter. To generate virus stocks, plasmids were linearized, in vitro transcribed using the mMESSAGE mMACHINE™ SP6 Transcription Kit (Thermo Fisher) and the resulting capped RNA electroporated into BHK-21 cells using a Gene Pulser Xcell electroporator (BioRad) and cultured for 48 h at 37 °C. The culture media was harvested and clarified by centrifugation. Virus stocks were aliquoted and stored at −80 °C.

### TK CHIKV-PV production and infection, and GCV treatment

5×10^6^ HEK293T/17 cells were seeded into a 10 cm dish 24 hours prior to transfection with p8.91, pCSTKW and the required viral envelope plasmid at a ratio of 1:1.5:1 µg [34, 35]. The plasmids were mixed with 1 mg/ml polyethylenimine (PEI; Merck) at a ratio of 1:6, left to incubate for 10-15 minutes and then added dropwise to the cells. The dish was then incubated at 37^°^C (5% CO_2_) for 48 hours before the supernatant was harvested and passed through a 0.45 µM filter. Following this the media was replaced and a second harvest undertaken a further 24 hours later. TK CHIKV-PV was titrated onto the target cells in a doubling dilution and GCV added to the cells (at stated concentrations) 24 hours later. The titre of virus required to kill ∼90% of the cells was determined 48 hours post GCV addition.

### CRISPR-CAS9 library preparation

We obtained the GeCKO v2 library from Addgene, amplified it by large scale electroporation with Endura competent cells (Lucigen) then amplified a sample by PCR to produce a library for next generation sequencing (NGS) on the Illumina MiSEq platform. The library passed the quality control checks for library representation.

We produced the Gecko library viruses by transfecting into HEK293T/17 cells with two lentiviral packaging vectors and harvesting the supernatant 2 days later. The lentivirus virions were titrated on MCF7 and HCC13965 cells using the functional readout of puromycin resistance. We generated a mutant cell pool from infecting 1.8 x10^8^ cells at a low MOI of 0.3 to ensure only 1 virus per cell. Cells were selected under puromycin 24 hours later and for 14 days total to produce a stable mutant pool then a T0 sample harvested from at least 6 x10^7^ cells.

The screen target population was then infected with CHKV-TK followed by GCV (75uM) 48 hours later, to achieve 90% of cell death in control cells. GCV was maintained in the media for 7 further days and dead cells removed by splitting and re-seeding. Surviving cells were then cultured for another 14 days and then harvested.

Genomic DNA was prepared for NGS as per the Gecko Nature methods paper, quality checked on a Bioanalyzer high sensitivity DNA chip. NGS was performed by BGI Genomics.

The next-generation sequencing data were processed with the Model-based Analysis of Genome-wide CRISPR/Cas9 Knockout (MAGeCK) algorithm [16].

### NGS and bioinformatic analysis

We used the MAGeCK-VISPR to analyse the CRISPR screening data [16]. Functional relationships between protein genes hits were detected using Sting analysis [36]. Next generation sequencing data from *Ceudovitox* screen has been deposited at GEO Datasets, under the accession number: GSE302317.

### Small molecule/compound and siRNA [PV and authentic virus]

Small molecules were applied at the indicated concentrations for 2 hours then CHIKV-luc added to the well. Luciferase activity was assayed 2 days later. Corresponding wells were analysed for cell toxicity with Cell Titre Glo reagent. For RNAi cells were incubated for 3 days after RNAi Max transfection (10nmol), then CHKV-luc applied and luciferase activity was assayed 2 days later. 1 × 10^5^ RLU/per well of CHIKV-luc virus was added for ‘high RLU’ challenge compared 2 × 10^4^ RLU/per well for low RLU challenge.

### Authentic (live) virus CHIKV entry inhibition assays

All tested compounds were added to BHK-21 cells 2 hours before infection, along with DMSO controls. For the assay, cells were seeded in 96 well plates at 2 × 10^4^ cells/well. CHIKV was then added to the cells at an MOI of 0.01 and 0.1 and fluorescence was measured every 6 hours over a 48-hour period. 6 technical replicates per treatment. Plates were imaged using the IncuCyte live cell imager (Sartorius) and images analyzed using the associated IncuCyte software (Sartorius).

## Acknowledgments

Thanks to the University of Sussex School of Life Sciences for their support.

## Supporting Information

***S1 Fig: Cytotoxicity of tested drugs at different concentrations***. *Cytotoxicity of different concentrations of (A) broad-spectrum MMP, Marimastat (purple line) and GM6001(green line) and (B) specific MMP targeting inhibitor drugs on A549. Specific MMP targeting drugs targeted ADAM10 (G1254023X, brown line), MMP13 (T26-C, purple line) and ADAM54 (Tinzapararin, orange line). Arbidol (red line) was used as positive control*.

## References

1. Roychoudhury S, Das A, Sengupta P, Dutta S, Roychoudhury S, Choudhury AP, et al. Viral Pandemics of the Last Four Decades: Pathophysiology, Health Impacts and Perspectives. International Journal of Environmental Research and Public Health. 2020 Dec 15;17(24). doi: 10.3390/ijerph17249411.

2. Marani M, Katul GG, Pan WK, Parolari AJ, Marani M, Katul GG, et al. Intensity and frequency of extreme novel epidemics. Proceedings of the National Academy of Sciences. 2021-08-31;118(35). doi: 10.1073/pnas.2105482118.

3. Afrough B, Eakins J, Durley-White S, Dowall S, Findlay-Wilson S, Graham V, et al. X-ray inactivation of RNA viruses without loss of biological characteristics. Scientific Reports 2020 10:1. 2020-12-08;10(1). doi: 10.1038/s41598-020-77972-5.

4. Bartholomeeusen K, Daniel M, LaBeaud DA, Gasque P, Peeling RW, Stephenson KE, et al. Chikungunya fever. Nature Reviews Disease Primers 2023 9:1. 2023-04-06;9(1). doi: 10.1038/s41572-023-00429-2.

5. Schilte C, Staikovsky F, Couderc T, Madec Y, Carpentier F, Kassab S, et al. Chikungunya Virus-associated Long-term Arthralgia: A 36-month Prospective Longitudinal Study. PLOS Neglected Tropical Diseases. 21 Mar 2013;7(3). doi: 10.1371/journal.pntd.0002137.

6. Zhang R, Kim AS, Fox JM, Nair S, Basore K, Klimstra WB, et al. Mxra8 is a receptor for multiple arthritogenic alphaviruses. Nature 2018 557:7706. 2018-05-16;557(7706). doi: 10.1038/s41586-018-0121-3.

7. Feng F, Bouma EM, Hu G, Zhu Y, Yu Y, Smit JM, et al. Colocalization of Chikungunya Virus with Its Receptor MXRA8 during Cell Attachment, Internalization, and Membrane Fusion. Journal of Virology. 2023-5-1;97(5). doi: 10.1128/jvi.01557-22.

8. Ottosen S, Parsley TB, Yang L, Zeh K, Doorn L-Jv, Veer Evd, et al. In Vitro Antiviral Activity and Preclinical and Clinical Resistance Profile of Miravirsen, a Novel Anti-Hepatitis C Virus Therapeutic Targeting the Human Factor miR-122. Antimicrobial Agents and Chemotherapy. 2014 Dec 23;59(1). doi: 10.1128/AAC.04220-14.

9. Gaur NK, Urankar S, Sengupta D, Chepuri VR, Makde RD, Kulkarni K. A cell based assay using virus-like particles to screen AM type mimics for SARS-CoV-2 neutralisation. Biochemical and Biophysical Research Communications. 2024/07/23;718. doi: 10.1016/j.bbrc.2024.150082.

10. Cantoni D, Wilkie C, Bentley EM, Mayora-Neto M, Wright E, Scott S, et al. Correlation between pseudotyped virus and authentic virus neutralisation assays, a systematic review and meta-analysis of the literature. Frontiers in Immunology. 2023 Sep 18;14. doi: 10.3389/fimmu.2023.1184362.

11. Tani H, Iha K, Shimojima M, Fukushi S, Taniguchi S, Yoshikawa T, et al. Analysis of Lujo Virus Cell Entry using Pseudotype Vesicular Stomatitis Virus. Journal of Virology. 2014-7-1;88(13). doi: 10.1128/jvi.00512-14.

12. Song H, Zhao Z, Chai Y, Jin X, Li C, Yuan F, et al. Molecular Basis of Arthritogenic Alphavirus Receptor MXRA8 Binding to Chikungunya Virus Envelope Protein. Cell. 2019/06/13;177(7). doi: 10.1016/j.cell.2019.04.008.

13. Yamamoto S, Suzuki S, Hoshino A, Akimoto M, Shimada T. Herpes simplex virus thymidine kinase/ganciclovir-mediated killing of tumor cell induces tumor-specific cytotoxic T cells in mice. Cancer Gene Ther. 1997;4(2):91–6. PubMed PMID: 9080117.

14. NE S, O S, F Z. Improved vectors and genome-wide libraries for CRISPR screening - PubMed. Nature methods. 2014 Aug;11(8). doi: 10.1038/nmeth.3047.

15. Shalem O, Sanjana NE, Hartenian E, Shi X, Scott DA, Mikkelsen TS, et al. Genome-Scale CRISPR-Cas9 Knockout Screening in Human Cells. Science. 2014-01-03;343(6166). doi: 10.1126/science.1247005.

16. Li W, Köster J, Xu H, Chen C-H, Xiao T, Liu JS, et al. Quality control, modeling, and visualization of CRISPR screens with MAGeCK-VISPR. Genome Biology 2015 16:1. 2015-12-16;16(1). doi: 10.1186/s13059-015-0843-6.

17. Szklarczyk D, Gable AL, Lyon D, Junge A, Wyder S, Huerta-Cepas J, et al. STRING v11: protein– protein association networks with increased coverage, supporting functional discovery in genome-wide experimental datasets. Nucleic Acids Research. 2018 Nov 22;47(Database issue). doi: 10.1093/nar/gky1131.

18. Henß L, Beck S, Weidner T, Biedenkopf N, Sliva K, Weber C, et al. Suramin is a potent inhibitor of Chikungunya and Ebola virus cell entry. Virology Journal. 2016 Aug 31;13(1). doi: 10.1186/s12985-016-0607-2.

19. Kaur P, Chu JJH. Chikungunya virus: an update on antiviral development and challenges. Drug Discovery Today. 2013/10/01;18(19-20). doi: 10.1016/j.drudis.2013.05.002.

20. Hu J, Van den Steen PE, Sang Q-XA, Opdenakker G, Hu J, Van den Steen PE, et al. Matrix metalloproteinase inhibitors as therapy for inflammatory and vascular diseases. Nature Reviews Drug Discovery 2007 6:6. 2007/06;6(6). doi: 10.1038/nrd2308.

21. Han J, Perez JT, Chen C, Li Y, Benitez A, Kandasamy M, et al. Genome-wide CRISPR/Cas9 Screen Identifies Host Factors Essential for Influenza Virus Replication. Cell Reports. 2018/04/10;23(2). doi: 10.1016/j.celrep.2018.03.045.

22. Shen D, Zhang G, Weng X, Liu R, Liu Z, Sheng X, et al. A genome-wide CRISPR/Cas9 knockout screen identifies TMEM239 as an important host factor in facilitating African swine fever virus entry into early endosomes. PLOS Pathogens. 2024 Jul 18;20(7). doi: 10.1371/journal.ppat.1012256.

23. Wang R, Simoneau CR, Kulsuptrakul J, Bouhaddou M, Travisano KA, Hayashi JM, et al. Genetic Screens Identify Host Factors for SARS-CoV-2 and Common Cold Coronaviruses. Cell. 2021/01/07;184(1). doi: 10.1016/j.cell.2020.12.004.

24. Thimmiraju SR, Kimata JT, Pollet J. Pseudoviruses, a safer toolbox for vaccine development against enveloped viruses. Expert Review of Vaccines. 2024-12-31;23(1). doi: 10.1080/14760584.2023.2299380.

25. Plesa G, McKenna PM, Schnell MJ, Eisenlohr LC. Immunogenicity of Cytopathic and Noncytopathic Viral Vectors. Journal of Virology. 2006 Jul;80(13). doi: 10.1128/JVI.00084-06.

26. Morizono K, Chen ISY. Receptors and tropisms of envelope viruses. Current opinion in virology. 2011 Jul 1;1(1). doi: 10.1016/j.coviro.2011.05.001.

27. Silva LA, Khomandiak S, Ashbrook AW, Weller R, Heise M, Morrison T, Dermody T. A Single-Amino-Acid Polymorphism in Chikungunya Virus E2 Glycoprotein Influences Glycosaminoglycan Utilization. Journal of Virology. 2013;88(5). doi: 10.1128/JVI.03116-13.

28. McAllister N, Liu Y, Silva LM, Lentscher AJ, Chai W, Wu N, et al. Chikungunya Virus Strains from Each Genetic Clade Bind Sulfated Glycosaminoglycans as Attachment Factors. Journal of Virology. 2020 Nov 23;94(24). doi: 10.1128/JVI.01500-20.

29. Schnierle BS, Schnierle BS. Cellular Attachment and Entry Factors for Chikungunya Virus. Viruses 2019, Vol 11, Page 1078. 2019-11-19;11(11). doi: 10.3390/v11111078.

30. Cabral-Pacheco GA, Garza-Veloz I, Rosa CC-Dl, Ramirez-Acuña JM, Perez-Romero BA, Guerrero-Rodriguez JF, et al. The Roles of Matrix Metalloproteinases and Their Inhibitors in Human Diseases. International Journal of Molecular Sciences. 2020 Dec 20;21(24). doi: 10.3390/ijms21249739.

31. Meléndez-Hevia E, Paz-Lugo Pd, Sánchez G, Meléndez-Hevia E, Paz-Lugo Pd, Sánchez G. Glycine can prevent and fight virus invasiveness by reinforcing the extracellular matrix. Journal of Functional Foods. 2021-01;76. doi: 10.1016/j.jff.2020.104318.

32. Dong S, Balaraman V, Kantor AM, Lin J, Grant DG, Held NL, Franz AWE. Chikungunya virus dissemination from the midgut of Aedes aegypti is associated with temporal basal lamina degradation during bloodmeal digestion. PLoS Neglected Tropical Diseases. 2017 Sep 29;11(9). doi: 10.1371/journal.pntd.0005976.

33. Wright E, Temperton NJ, Marston DA, McElhinney LM, Fooks AR, Weiss RA, et al. Investigating antibody neutralization of lyssaviruses using lentiviral pseudotypes: a cross-species comparison. Journal of General Virology. 2008/09/01;89(9). doi: 10.1099/vir.0.2008/000349-0.

34. Zufferey R, Nagy D, Mandel RJ, Naldini L, Trono D, Zufferey R, et al. Multiply attenuated lentiviral vector achieves efficient gene delivery in vivo. Nature Biotechnology 1997 15:9. 1997/09;15(9). doi: 10.1038/nbt0997-871.

35. Mather ST, Wright E, Scott SD, Temperton NJ. Lyophilisation of influenza, rabies and Marburg lentiviral pseudotype viruses for the development and distribution of a neutralisation - assay-based diagnostic kit. Journal of Virological Methods. 2014/12/15;210. doi: 10.1016/j.jviromet.2014.09.021.

36. Szklarczyk D, Kirsch R, Koutrouli M, Nastou K, Mehryary F, Hachilif R, et al. The STRING database in 2023: protein–protein association networks and functional enrichment analyses for any sequenced genome of interest. Nucleic Acids Research. 2022 Nov 12;51(D1). doi: 10.1093/nar/gkac1000.

